# Intraspecific variation alters ecosystem and next-generation performance as much as temperature

**DOI:** 10.1101/332619

**Authors:** Allan Raffard, Julien Cucherousset, Frédéric Santoul, Lucie Di Gesu, Simon Blanchet

## Abstract

Phenotypic or genotypic variation within species affects ecological processes, from populations to ecosystems. However, whether the ecological imprint of intraspecific variation is substantial compared to key environmental drivers, and persistent enough to carry over to next generations is still questioned. Here, we experimentally showed that intraspecific variation manipulated in a freshwater fish (the European minnow, *Phoxinus phoxinus*) led to ecological and transgenerational carry-over effects that were as strong as those of varying temperature by 2°C. Specifically, variation in fish body mass, growth rate and activity altered the size and abundance of prey, which ultimately affected traits and survival of the next fish generation. Temperature variation modulated other ecosystem functions (e.g. litter decomposition) that were also associated to transgenerational carry-over effects. Our results demonstrate that shifting genotypes or phenotypes in wild populations can have substantial and persistent consequences on ecosystems with a similar intensity than climatic variation.

## Introduction

Investigating species trait variability provides a mechanistic framework to predict the links between biodiversity and ecosystem functions (Violle et al. 2007). While trait variation has primarily been quantified at the species level, intraspecific trait variation accounts for a high proportion of the total trait variability in communities (Zhao et al. 2019). Furthermore, intraspecific trait variation has been shown to alter community structure and ecosystem functioning (Rudolf and Rasmussen 2013, Hendry 2016), indicating that intraspecific variation is a significant driver and predictor of ecological dynamics (Des Roches et al. 2018, Raffard et al. 2019b).

Intraspecific trait-mediated ecological effects can sometimes be strong and persist long enough to affect the subsequent generations through ‘inheritance’ of the surrounding environment (Odling-Smee et al. 2013). Indeed, next generations can experience environmental conditions that have been modified by previous generations, subsequently affecting next-generations performance such as foraging efficiency, growth and survival (Matthews et al. 2016, Best et al. 2017). These “transgenerational carry-over effects” can – under some circumstances – affect patterns of selection and hence the evolutionary dynamics of species (i.e. eco-evolutionary feedbacks, Schoener 2011, Hendry 2016). While these ecological and transgenerational carry-over effects have been the focus of recent experimental and theoretical works (Matthews et al. 2016, Brunner et al. 2017), whether intraspecific variation really matters for the long-term dynamics of complex ecosystems is still questioned (Thompson 1998, Schoener 2011).

This issue can be addressed by comparing the intensity of the effects induced by intraspecific variation to those induced by key environmental factors affecting both ecological and evolutionary dynamics, such as temperature (Schoener 2011). Intraspecific variation can affect ecological processes with an intensity similar to that of environmental drivers (El-Sabaawi et al. 2015). However, whether the ecological imprints of intraspecific variation are strong and persistent enough to affect subsequent generations, and whether these transgenerational carry-over effects are similar in strength to those of indisputably important environmental drivers is still unknown. Addressing this question is an important step for determining the relative contribution of intraspecific variation in predicting the responses of ecosystems to global change.

Here, we compared the consequences of intraspecific variation on ecosystem properties and on the performance of subsequent generations to those induced by changes in temperature, and assessed which of a series of traits contribute the most to these consequences. Temperature is a key abiotic factor directly affecting ecosystem functions such as primary productivity (Yvon-Durocher et al. 2015), and imposing a strong selective pressure on organism traits (Brown et al. 2004, Rey et al. 2016). We ran a two-phase experiment (Matthews et al. 2011, 2016) (Figure 1) manipulating (i) trait variation in a freshwater fish (European minnow, *Phoxinus phoxinus*) by selecting individuals from populations having evolved distinct genotypes and phenotypes (Figure S1 and S2, see also Raffard et al. 2019a) and (ii) water temperature by setting mesocosms to differ by 2°C throughout the experiment (Figure S3). An increase in temperature of 2°C represents the general warming expectations for freshwaters over the next 40 years (IPCC 2014). During the first experimental phase (*ecological effects*), we compared the strength of the effects of trait variation among adult minnows to the strength of the effects of temperature variation on prey community structure and ecosystem functioning (Figure 1). Adults were then removed from the mesocosms and replaced by juveniles with a common origin for the second experimental phase (*transgenerational carry-over effects*). We tested how the ecological variations induced during the first phase (due to intraspecific and/or temperature variation) affected the performance of juveniles.

**Figure 1.**
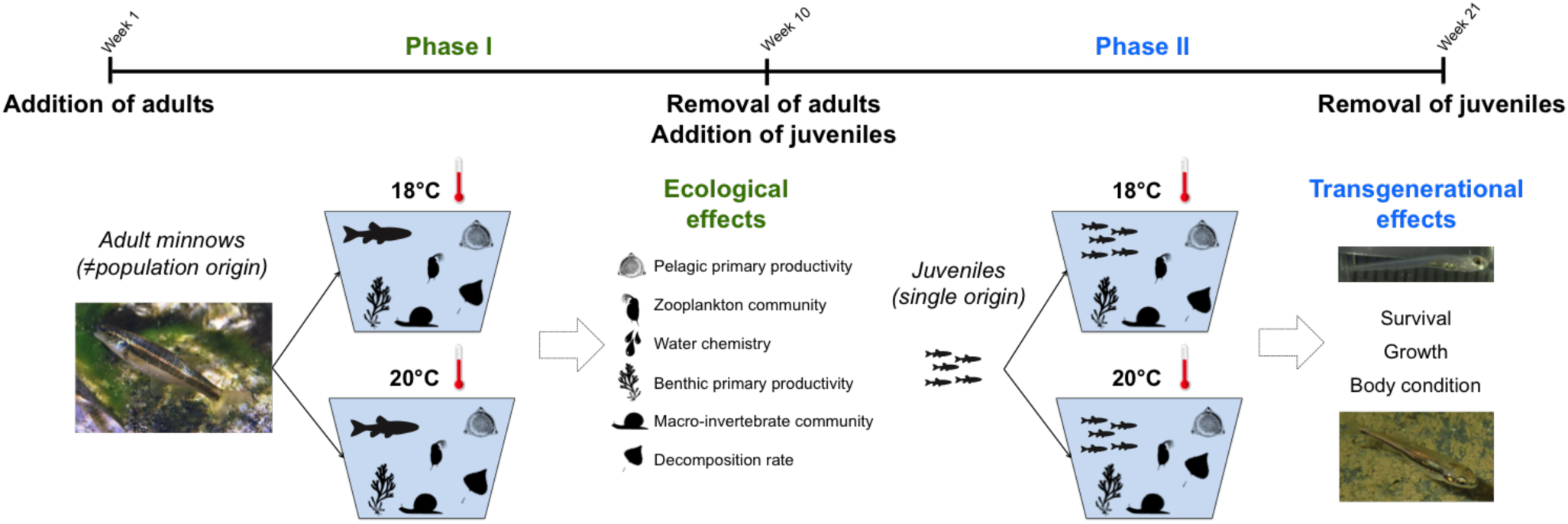
Experimental design used to test the ecological (phase 1) and transgenerational carry-over (phase 2) effects of intraspecific variation among adult minnows (*Phoxinus phoxinus*) and temperature variation.

## Material and Methods

### Study species

European minnow (*Phoxinus phoxinus*) was used as the model species. *P. phoxinus* is a small-bodied (maximum length: ~80 mm, mean generation time: ~2 years) cyprinid fish species widely distributed in Western Europe. *P. phoxinus* lives in relatively cold lentic and lotic waters. It is a generalist species that feeds on small invertebrates, algae and zooplankton.

In September 2016, we collected adult minnows by electrofishing in six rivers in southwestern France (Figure S1). We selected populations that were isolated geographically and had distinct environments (Figure S4) to favor both genetic and phenotypic divergences among populations. Accordingly, the mean genetic divergence among populations was F_st_ = 0.162 (measured using 17 microsatellites, min-max = 0.043-0.313). The sampled populations also varied for important functional traits, such as body mass and metabolic rate (Figure S2). Furthermore, we have previously shown that minnow populations in this river system were characterised by unique “functional syndromes” (set of trait variation and covariation) that resulted from a combination of random drift and adaptation to environmental factors (Raffard et al. 2019a). All fish collections and husbandry for adults and juveniles were conducted in accordance with sampling permits obtained from local authorities (25-08-2016, 24-05-2016, 09-273, SA-013-PB-092, A09-3). Fish from different populations were reared at a similar density and separately for ~6 months in 1100 L outdoor tanks to minimize prior environmental effects on phenotypes. During rearing, the fish were fed ad-libitum with a mixture of pelletized food and dead chironomids.

### Phase 1: effects of intraspecific variation and temperature on ecological processes

The experiment consisted of 72 replicated mesocosms placed in a greenhouse with a 12:12 h light-dark photoperiod. Mesocosms were filled with 100 L of tap water and 1 cm of gravel covering the bottom of each tank. Tanks were covered with a 1 cm plastic mesh net to prevent fish escapes. Nutrients were added to the mesocosms using 5 mL of solution containing nitrogen and phosphorus (ratio N: P: K = 3.3: 1.1: 5.8) on December 2^nd^ 2016. Each mesocosm was then inoculated with 200 mL of a concentrated solution of phytoplankton from a unique lake origin (Lake Lamartine, France 43°30’21.5”N, 1°20’32.7”E) on December 12^th^ 2016. Two months later (February 15^th^ 2017), an additional 200 mL of concentrated solution of zooplankton from the same lake was added to each mesocosm. Finally, we inoculated each mesocosm with sediment and macroinvertebrates (i.e., mainly Gastropoda and Bivalvia) from Lake Lamartine.

Each tank was assigned to one of twelve treatments (each replicated six times) according to a full-factorial design with intraspecific variation (i.e., population origin, six levels corresponding to each population) and temperature (two levels: low and high temperature) as the main factors (Figure 1). Water temperature was controlled and adjusted using a *Blue Marine®* water chiller and a stainless steel coil placed in each tank through which a flux of water (independent from the water of the tanks) flowed at either 18°C or 21°C. On average, the low and high water temperature treatments differed by 2.08°C according to natural seasonal variations (Figure S3).

In March 2017, adult fish were weighed to the nearest 0.01 g and a single fish was introduced to each mesocosm. This individual-based approach prevented the experimental ecosystems from collapsing due to the over-density of top consumers and allowed the ecological effects of individual trait variation to be measured. After 73 days (Figure 1), each fish was removed and phenotyped. We measured six traits on each individual including body mass, growth rate, metabolic rate, activity, aspect ratio and gut length (see supplementary method for details). Concomitantly, we measured multiple community and ecosystem parameters to evaluate differences in ecological processes among treatments: Pelagic and benthic algae stocks, filamentous algae, decomposition rate, abiotic parameters, Copepoda and Cladocera abundances and size, and macroinvertebrates (Bivalvia and Gastropoda) abundance (see supplementary method for details).

### Phase 2: transgenerational effects of ecological differences on juvenile performance

After the removal of adult fish on June 13^th^ 2017, 45 juvenile minnows were introduced to each mesocosm. We used juveniles from a single origin (i.e., fish farm, *Amorvif EURL*) to control for potential genetic effects. Juveniles were introduced as soon as possible after hatching to increase the possibility of differential mortality and/or ontogenetic plasticity. Therefore, juveniles were introduced when they were only two weeks old as stage III larvae (Figure S5). They were not manipulated (i.e., weighted and/or measured) before being randomly introduced in the mesocosms to limit potential mortality. The juveniles were removed from the mesocosms 79 days later, and we measured several proxies for their fitness (i.e. performance). Individuals were counted to assess survival, weighed to the nearest 0.001 g to assess growth rate (assuming all juveniles had the same initial body mass, we used the final body mass of juveniles as a measure of growth rate), and measured in length to the nearest 0.1 mm (using ImageJ) to assess the body condition, which was calculated as the residuals of the relationship between individual body mass and length.

### Statistical analyses

Two adult individuals died before the end of phase 1, so we discarded these two replicates from the analyses. Moreover, we identified six tanks in which crayfish had been inadvertently introduced; we discarded these six replicates because crayfish are known to have disproportionally strong impacts on ecosystems (Alp et al. 2016). As such, the final analyses were run on 64 replicates.

First, we compared the magnitude of the effects of intraspecific variation and temperature increase on ecological (phase 1) and evolutionary (phase 2) dynamics. To do so, we used a meta-analytic approach consisting of first running linear models linking each ecological or evolutionary parameter (dependent variables) to the explicative variables, i.e., intraspecific variation (categorical factor, six levels), temperature (categorical factor, two levels) and the resulting two-term interaction. The interaction term was removed when not significant because it prevents the interpretation of simple terms (Nakagawa and Cuthill 2007). From these linear models, we calculated the standardized effect sizes eta squared (Levine and Hullett 2002) (*η*^2^) as follows: *η*^*2*^ = *SS*_*x*_*/SS*_*tot*_, where *SS*_*x*_ is the sum of squares for the effect of interest (intraspecific variation, temperature or the interaction term, if significant) and *SS*_*tot*_ is the total sum of squares. Sums of squares were extracted from type II analysis of variance when the interaction was not in the model and from type III analysis of variance when the interaction was significant (Langsrud 2003). Finally, the mean effect size (MES) values of intraspecific variation and temperature across the ecological or evolutionary parameters were compared using t-test.

Next, we assessed which individual trait contributed the most to ecological differentiations and subsequently to carry-over effects to the next generation using a causal analysis. Since we aimed at identifying the mechanisms by which the mesocosms diverged, we included the six traits measured on adult fish from phase 1 because these traits can drive ecological processes (Hildrew et al. 2007). We used path analyses (Grace 2006) to set a full model based on biologically rational paths and the visual inspection of the variance-covariance matrix, and all variables were scaled to the mean to facilitate the comparison. We first ran a model linking individual traits and temperature to ecological parameters. A full model was constructed and simplified by removing sequentially weak and/or nonsignificant paths until reaching a model that was correct statistically (i.e., a model that best fit the observed covariance matrix based on the maximum likelihood χ^2^ statistic (Grace 2006), while leading to the lowest Akaike Information Criteria (AIC) value. We finally extracted the absolute values of path coefficients from the final model to tease apart the direct and indirect effects of individual traits and temperature on the ecological parameters. We then ran a second path-analysis linking ecological parameters to performance parameters of juveniles, while fixing the links found in the first model (i.e. traits to ecological parameters) so as not to over-fit the model. Statistical analyses were performed using R software (R Core Team 2013), and path analyses were run using Amos (Arbuckle 2014).

## Results

In the first phase (Figure 1), we found that the effects of intraspecific variation in adult minnows on ecosystem properties (measured over all ecological parameters) had a similar intensity than those induced by temperature increase (mean effect size (MES) ± standard error = 0.103 ± 0.018 and MES ± SE = 0.078 ± 0.036 for intraspecific variation and temperature, respectively; *t* = 0.624, d.f = 18, *p* = 0.540, Figure 2). Nonetheless, the effects differed between ecological parameters (Figure 2b, Figure S6). For instance, intraspecific variation had the strongest effect on abundance and size of the fish prey community (Cladocera, Copepoda and Gastropoda), whereas increasing temperature had a particularly strong effect on decomposition rate (Figure 2b, see also Figure S7, S8 and Table S1 for details). The interaction term between temperature and intraspecific variation was significant only for benthic algae stock (*F* = 10.831, d.f = 5,52, *p* = 0.022), indicating that the ecological effects of intraspecific variation were temperature-independent for most ecological parameters.

**Figure 2.**
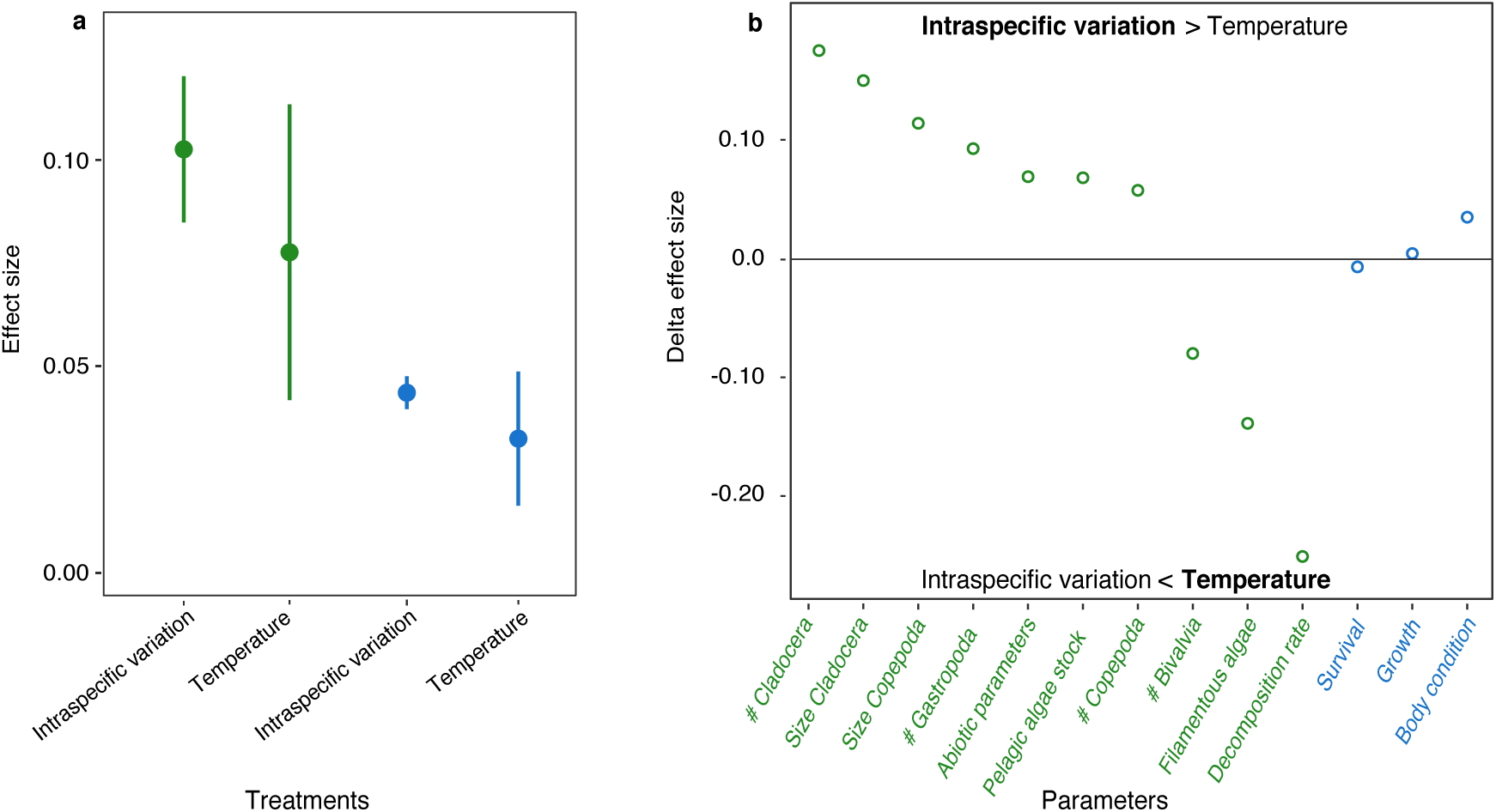
(a) Eta squared (η^2^) effect size of the intraspecific variation among adults and temperature on ecological (green) and performance (blue) parameters. Error bars represent ± 1 SE. (b) Delta of effect sizes (η^2^) of intraspecific variation and temperature on ecological and performance parameters. Positive values indicate a higher effect of intraspecific variation, and negative values indicate a higher effect of temperature.

We found that intraspecific variation in fish body mass and activity were the most influential functional traits and that variation in each of these two traits had similar –although weaker-ecological effects than temperature variation (Figure 3a). We further found that trait variation acted mainly directly on ecological parameters, and minor indirect effects were detected (Figure 3a). For instance, adult body mass positively affected the abundance of Copepoda directly, subsequently leading to an indirect effect on the abundance of Cladocera (Table S2). The ecological effects of temperature were both direct and indirect (Figure 3a). For instance, temperature directly increased Bivalvia abundance, which positively affected the abundance of Copepoda and the size of Cladocera, hence representing an indirect effect of temperature on the zooplankton community (Table S2). The covariance structure of the simplified path model did not differ from that of the data (χ^2^ = 87.064, d.f = 114, p = 0.971), indicating that the data were well supported by the model.

**Figure 3.**
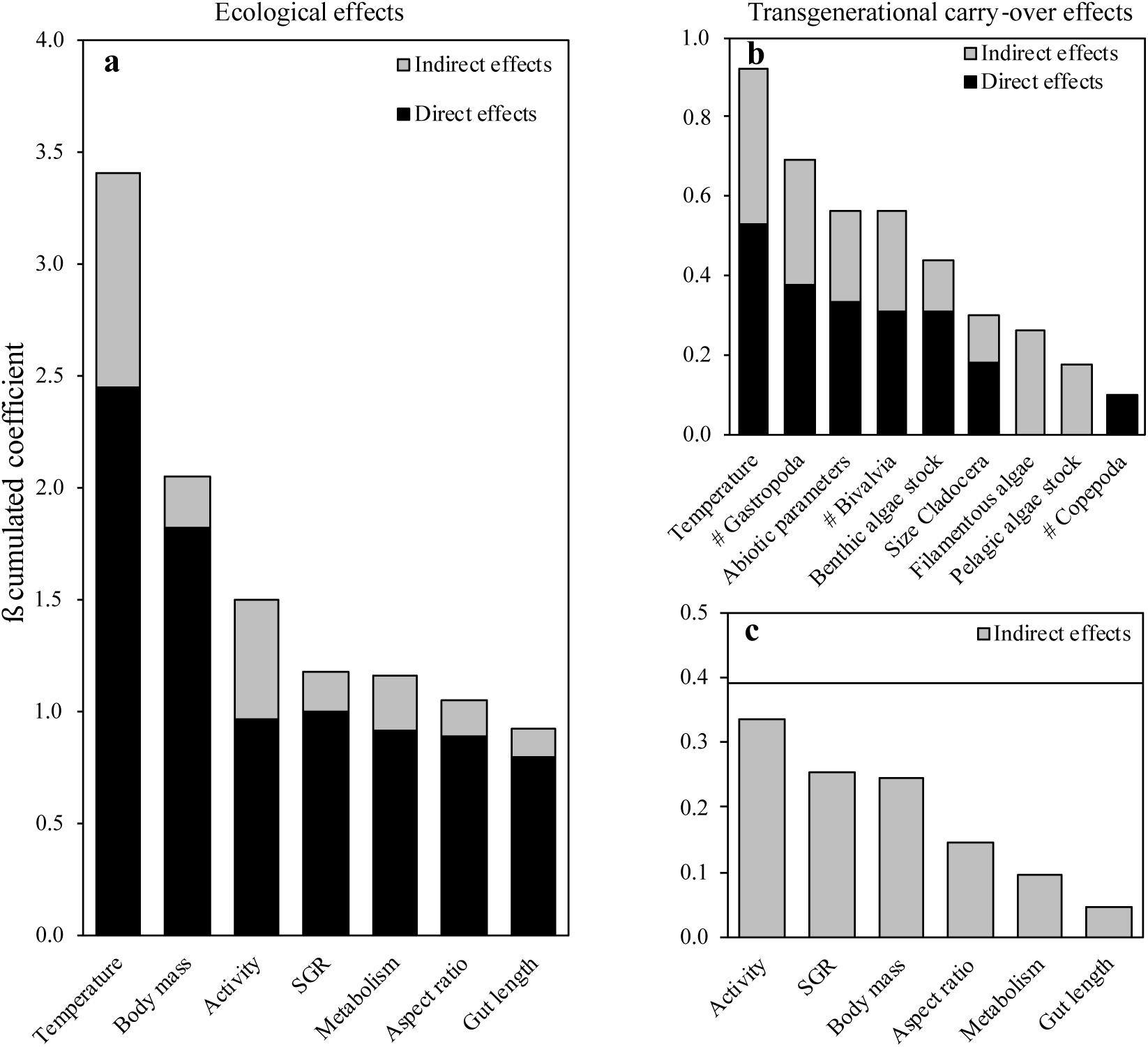
Cumulated absolute β path coefficients extracted from the simplified path-analysis described the direct and indirect relationships between intraspecific trait variation among adult minnows, temperature, ecological parameters (a) and performance parameters (b) and (c). The effects of intraspecific trait variation on performance of juveniles can only be indirect because the adult fish were removed before phase 2 of the experiment. The horizontal bar in (c) represents the indirect effects of temperature on performance parameters. The abbreviation ‘SGR’ stands for specific growth rate of adult minnows, and “#” for organism abundance.

In the second phase (Figure 1), we found that the strength of the effect sizes of intraspecific variation and temperature on the performance of juveniles (survival, growth rate and body condition) were similar (MES ± SE = 0.044 ± 0.004 and MES ± SE = 0.032 ± 0.016 for intraspecific variation and temperature, respectively, *t* = 0.665, d.f = 4, *p* = 0.542, Figure 2). The average effect sizes of intraspecific and temperature variations on the performance of juveniles were half the intensity of those on ecological parameters (Figure 2), indicating that transgenerational carry-over effects induced by the initial treatments were weaker in average than ecological effects observed in the first phase. Nonetheless, we found evidence for transgenerational carry-over effects of intraspecific variation since juvenile survival, growth rate and body condition were influenced by ecological parameters (e.g. benthic algae stock, Copepoda and Gastropoda abundance) that were themselves controlled by intraspecific variation (Figure 3b, Table S3). Amongst the adult traits, activity, growth rate and body mass contributed the most to the carry-over effects on juvenile performances (Figure 3c). For instance, active individuals decreased the abundance of Gastropoda, which subsequently affected juvenile survival and growth rate (Tables S2 and S3). Temperature increase mainly impacted -negatively-juvenile survival (i.e., survival increased as temperature decreased, Table S3). Also, juvenile survival was related to juvenile growth rate (density-dependent growth rate), leading to an indirect effect of temperature on juvenile growth rate (Table S3). Finally, juvenile body condition covaried with both juvenile survival and growth rate, and was affected by multiple ecological parameters (Table S3) which were directly affected by trait variation and temperature (Table S2). The covariance structure of the simplified path model for the phase 2 did not differ from that of the data (χ^2^ = 148.875, d.f = 196, p = 0.995), indicating that the data were well supported by the model.

## Discussion

We found that the ecological consequences of intraspecific and temperature variations were similar in strength but acted independently on different ecological parameters. Adult minnows from distinct populations modulated both the abundance and the size of their prey (zooplankton and Gastropoda), and these effects were predicted by variation among individuals in functional traits. In contrast, temperature variation strongly accelerated leaf litter decomposition, probably because warmer temperatures stimulate bacterial activity. These ecological imprints led to transgenerational carry-over effects on the performance of the next generation, which were also similar in strength between intraspecific variation and temperature. For instance, the survival of juveniles was higher in the low-temperature than in the high-temperature treatment, whereas the growth rate of juveniles differed depending on the trait characteristics of adult minnows introduced at the onset of the experiment. Overall, our results demonstrate that intraspecific variation can substantially affect ecosystem properties, which fosters ecological legacy to the next generations as much as temperature does.

Using a fine phenotypic screening of individual functional traits, we identified the traits contributing the most to the ecological effects of intraspecific variation. We revealed that multiple traits mediated the top-down control of minnows on prey community structure, ecosystem processes and subsequent transgenerational carry-over effects. Some of these traits were different among populations (e.g. body mass, metabolic rate), and both genetic (adaptive or not) and environmental factors underlined these trait differences among these populations (Raffard et al. 2019a). Body mass and metabolic rate are functional traits affecting the energetic needs and assimilation of individuals (Brown et al. 2004, Hildrew et al. 2007), leading to trophic differences among individuals. Our results demonstrate that population differences on these traits can strongly contribute to subsequent ecosystem differentiations. We further identified traits, such as activity, that mainly varied within populations but that also contributed to changes in ecosystem functioning. Behavioural variation in populations allows individuals to increase their performance through niche partitioning, ultimately affecting communities and ecosystems (e.g. Gastropoda abundance, Wolf and Weissing 2012). This confirms that individuals are not functionally equal neither among nor within populations (Wolf and Weissing 2012, Royauté and Pruitt 2015), and that –as recently shown in another fish species (Schmid et al. 2019)- it is possible to use these trait values to predict changes in important ecological properties.

Furthermore, we provide novel insights by demonstrating that these traits can also generate substantial transgenerational carry-over effects. The carry-over effects that we detected on subsequent generations are important as they suggest that the ecological effects of intraspecific variation in minnow are strong and long-lasting enough to modulate the whole dynamics of ecosystems. These effects correspond to indirect effects of trait variation among adult minnows on the performance of juveniles, which were mediated by the direct consequences of adult minnows on the ecological theatre. Multiple ecological parameters affected the performance of juveniles, such as the resource of juveniles (e.g. size of Cladocera) or the abiotic composition of their environment (e.g. pH or oxygen). Unsurprisingly, adult traits that have the highest ecological effects foster the highest transgenerational carry-over effects on juvenile performance, meaning that variation in these traits can be used to predict both direct ecological effects and indirect transgenerational effects of intraspecific variation. Currently, very few studies have demonstrated the existence of transgenerational carry-over effects, and most of them have focused on model organisms (e.g. Turcotte et al. 2013, Matthews et al. 2016). Our study extends the taxonomic scope of transgenerational effects, and suggests that this process does not concern only species with strong evolutionary divergences (Matthews et al. 2011).

Interestingly, intraspecific and temperature variations acted additively but not interactively on ecosystem properties and on the performance of next generation. Indeed, we identified only one significant interaction term between intraspecific variation and temperature on benthic algae stock. This finding confirms that the ecological consequences of intraspecific variation are often independent from the abiotic context (El-Sabaawi et al. 2015), which also seems to be the case for the transgenerational carry-over effects induced by intraspecific variation. This independence is surprising, since local adaptation for specific fitness traits and/or for reaction norms often leads to strong context dependency in the responses of organisms to local abiotic conditions (Kawecki and Ebert 2004), and we may have observed cascading interactive effects of intraspecific variation on ecological and evolutionary dynamics (Rosenblatt and Schmitz 2016). This finding is important because the absence of strong interactive effects reduces biological complexities and may therefore improve our ability to forecast the ecological and evolutionary consequences of environmental and biodiversity changes (Beckage et al. 2011).

In conclusion, we demonstrated that the magnitude of the ecological and transgenerational carry-over effects of intraspecific variation were as strong as the effects of temperature variation. This finding strongly supports the growing view that intraspecific variation and resulting transgenerational carry-over effects are not random noise, but are rather biologically substantial. Current environmental changes are rapid and can directly affect ecosystem functioning (Yvon-Durocher et al. 2015). These changes can also directly modulate the distribution of intraspecific trait variation in landscapes and thereby indirectly affect the dynamics of biological systems (Matthews et al. 2016, Brunner et al. 2017). Importantly, we demonstrate that some specific traits that are often targeted by humans (body size and activity, Biro and Post 2008) can be used to forecast ecosystem modifications under current environmental changes. To sum up, our results reinforce recent reports that changes and shifts in intraspecific variation of wild populations could be as harmful as considerable environmental changes (e.g., warming) to biological dynamics, and that this facet of biodiversity should therefore be conserved adequately (Mimura et al. 2016).

## Acknowledgements

We warmly thank Jose M. Montoya, Jean Clobert, Michel Loreau, Delphine Legrand and Jérôme G. Prunier for their valuable comments. We thank Lucas Mignien, Kéoni Saint-Pe and Yoann Buoro for their help during the experimental work. AR was financially supported by a doctoral scholarship from the Université Fédérale de Toulouse. This work was undertaken at SETE and EDB, which are part of the “Laboratoire d’Excellence” (LABEX) entitled TULIP (ANR-10-LABX-41).

## Authors’ contributions

A.R and S.B conceived the study. A.R and L.DG carried out the experiment with contributions from S.B and J.C. A.R performed the statistical analyses. A.R, S.B, J.C and F.S interpreted and discussed the results. A.R, S.B and J.C wrote the article, and all authors made corrections.

