## supplementary information for "Intraspecific variation alters ecosystem and next-generation performance as much as temperature"

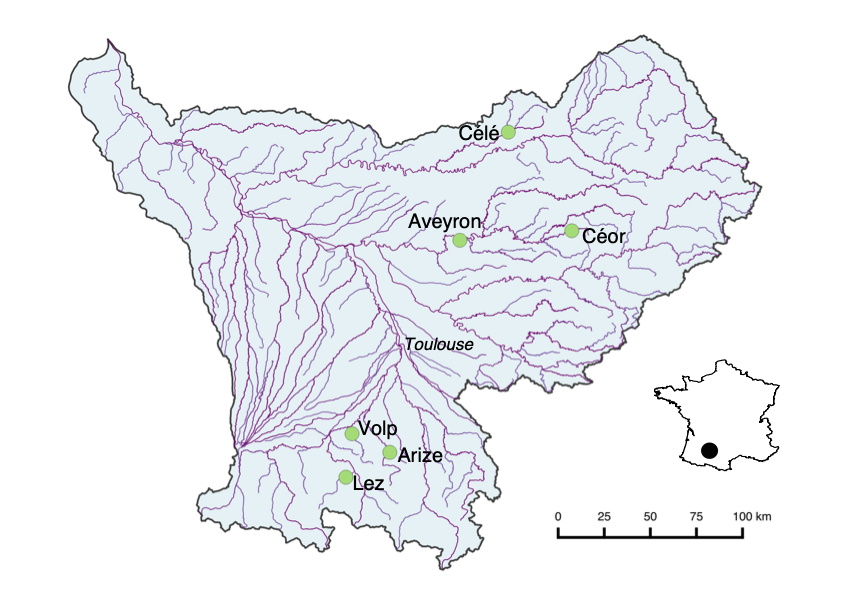

**Fig. S1.** Location of the six populations of adult minnows (*Phoxinus phoxinus*).

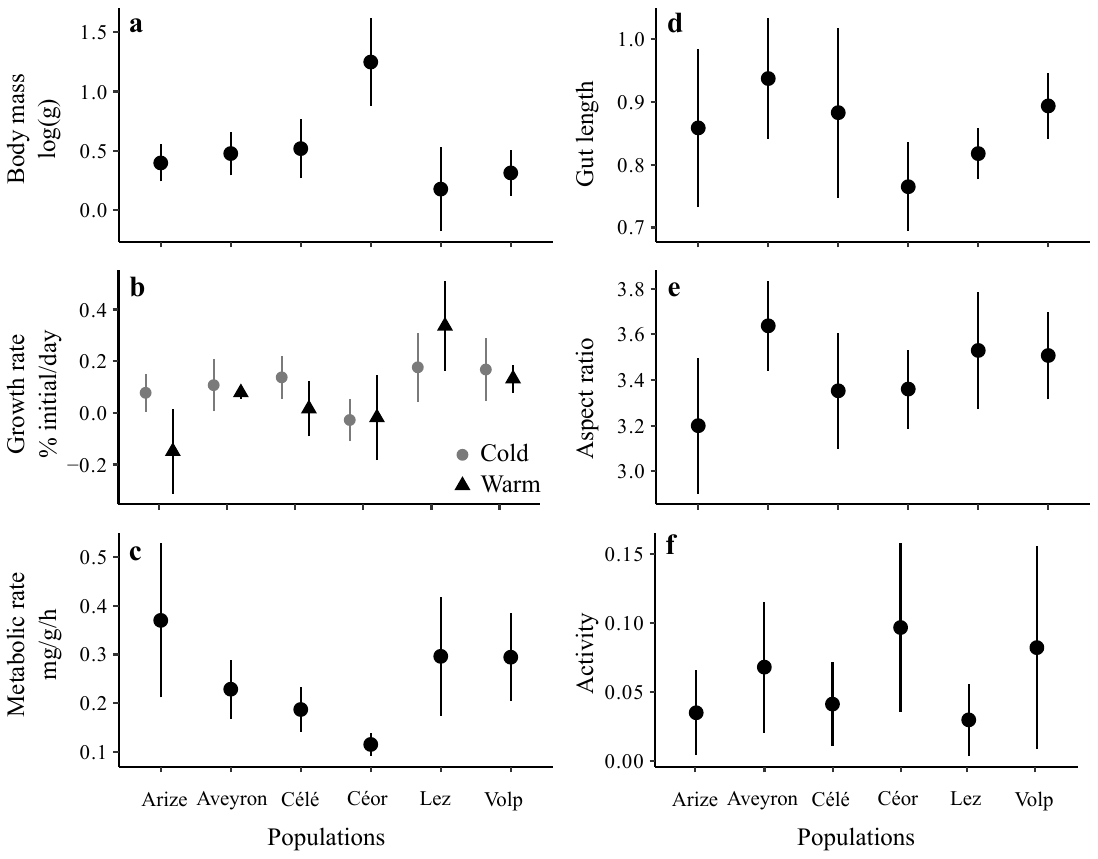

**Fig. S2. Phenotypic variation among populations of European minnows. (a)** Mean body mass (log-transformed), **(b)** growth rate (% of initial mass gain), **(c)** metabolic rate (i.e. oxygen consumption in mg.g^-1^.h^-1^), **(d)** gut length (ratio), **(e)** aspect ratio (ratio) and **(f)** activity (frequency of daily activity). Body mass was measured at the onset of the experiment and was significantly different among populations (*F* = 7.404, d.f = 5, 58, *p* < 0.001). Morphological traits (gut length and aspect ratio) were measured at the end, while growth rate and activity were measured during the experiment. Adult growth rate depended upon the interaction between their origin and temperature (*F* = 4.230, d.f = 5, 51, *p* = 0.002). Metabolic rate was significantly different among populations (*F* = 5.473, d.f = 5, 54, *p* < 0.001), gut length tended to be different among populations (F = 2.159, d.f = 5, 54, *p* = 0.072), and aspect ratio and activity were not different among populations (*p* > 0.1). Temperature did not affect these traits. Error bars represent ± 1 SE.

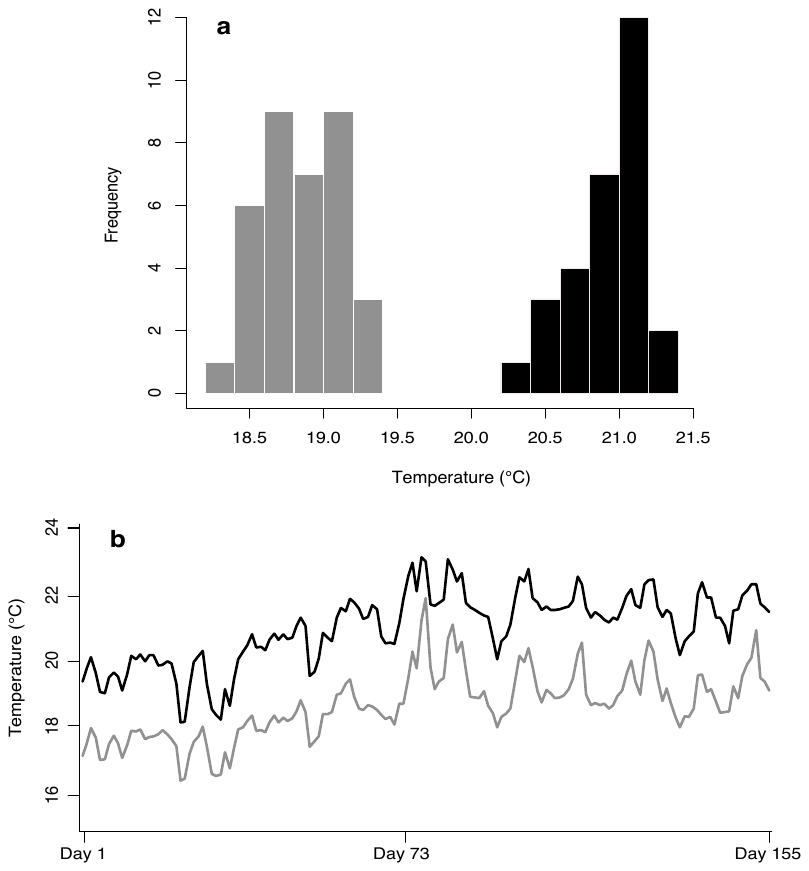

**Fig. S3. (a)** Frequency distribution of mean water temperature measured in each mesocosm from the low and high temperature treatment. Mean water temperature of mesocosms was significantly different (*t* = -32.647, d.f = 62, *p* < 0.001) between the low (grey, mean ± SE = 18.83 ± 0.04) and the high (black, mean ± SE = 20.91 ± 0.05) temperature treatments. **(b)** Averaged daily water temperature of two randomly chosen mesocosms among the two temperature treatments (low and high temperature treatments in grey and black, respectively). Water temperature from each tank was continuously recorded during the experiment using automatic data loggers (Hobo®).

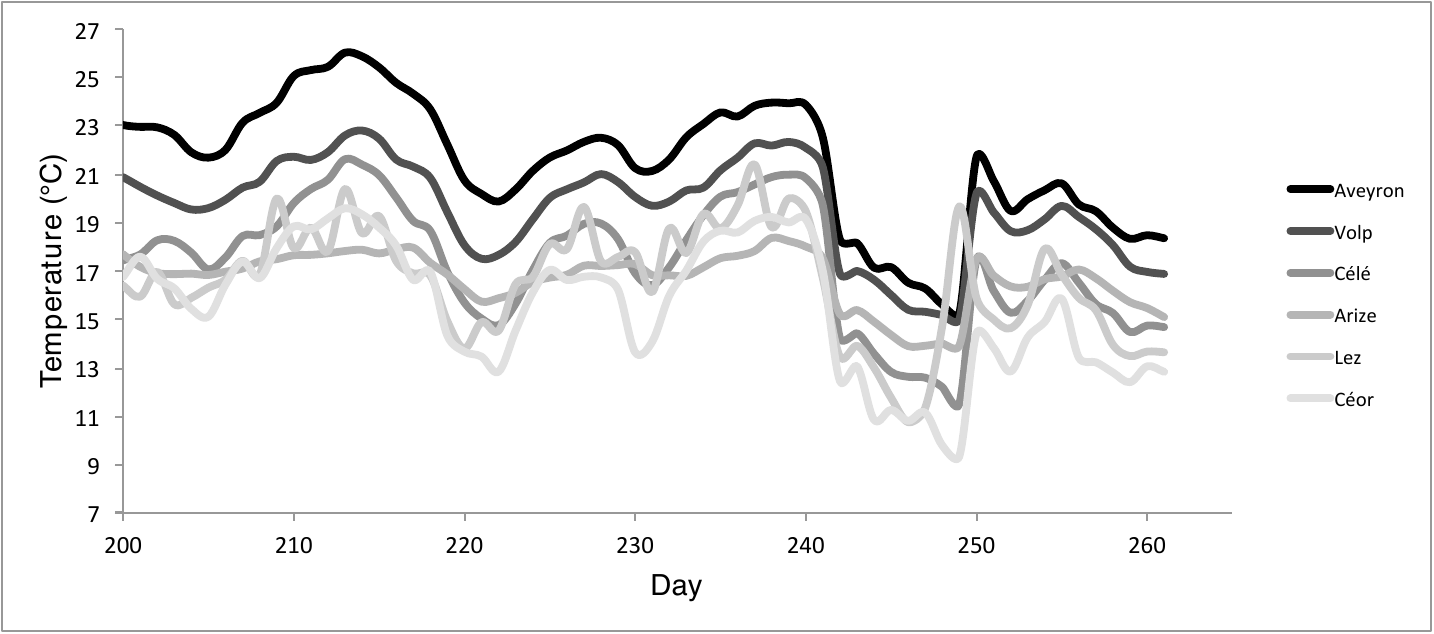

**Fig. S4.** Daily water temperature in each of the six rivers where adult minnows (*Phoxinus phoxinus*) were collected for phase 1 of the experiment. Water temperature was measured continuously in each river at the section where fish were sampled from the 21^st^ of July to the 12^th^ of September 2017 (the growing season) using automatic recorders (Hobo ®).

**Fig. S5.** Two weeks old juvenile minnows (*Phoxinus phoxinus,* stage III larvae, Pinder 2001) at the start of the phase 2 **(a)** and after 11 weeks at the end of the experiment **(b)**.

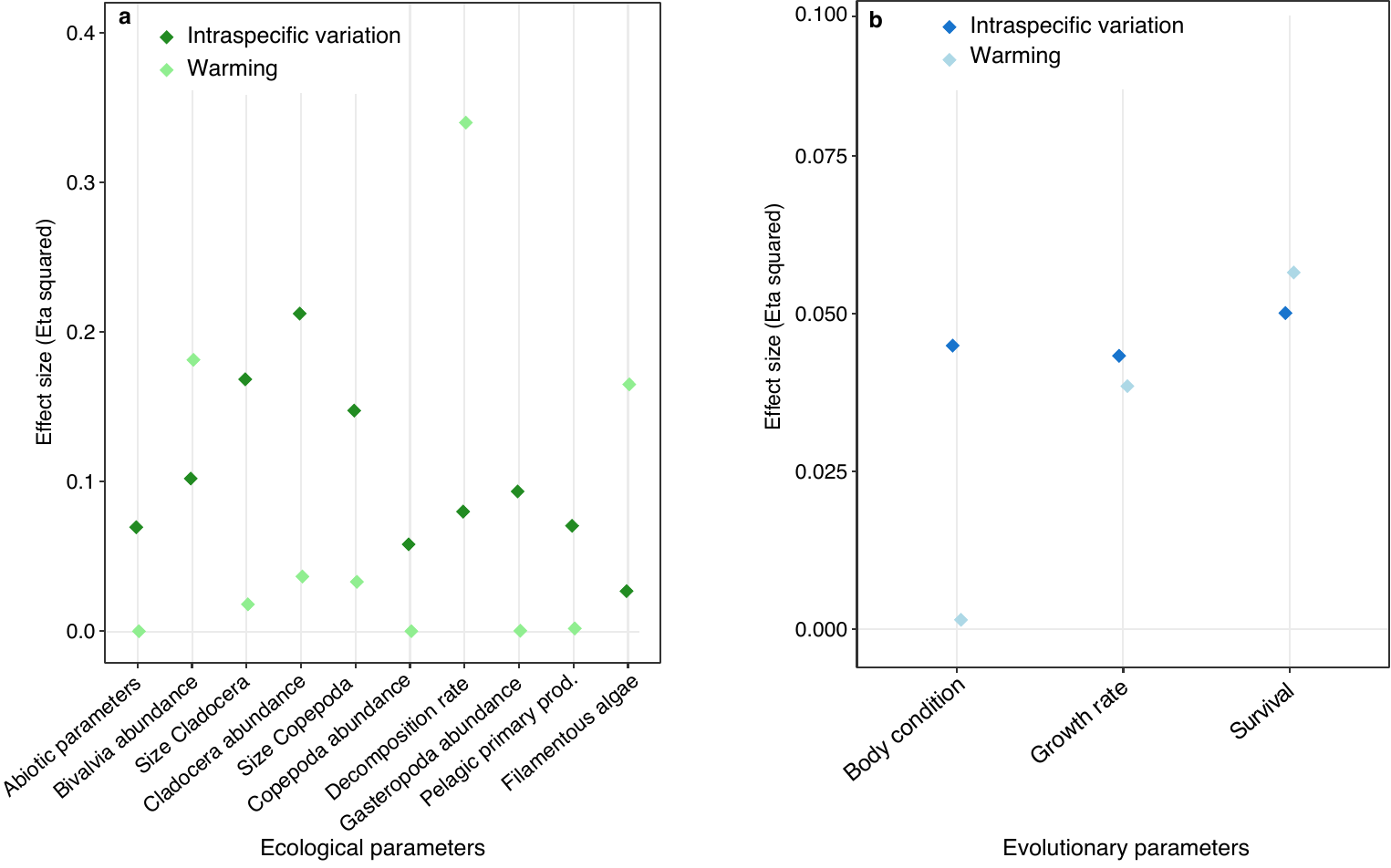

**Fig. S6.** Effect size (Eta squared) of intraspecific variation and warming on ecological **(a)** and performance **(b)** parameters.

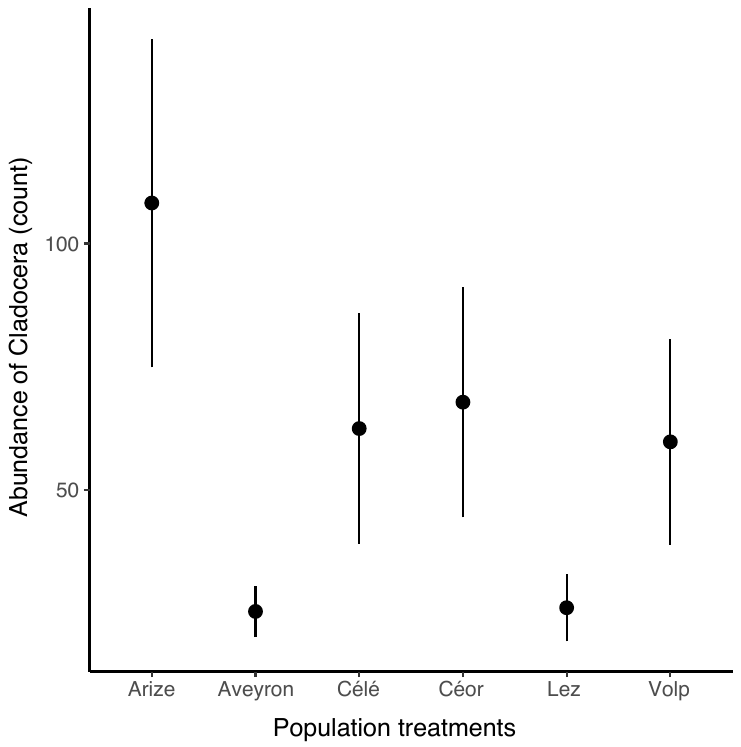

**Fig. S7.** Abundance of Cladocera (count data) remaining in the mesocosms at the end of the phase 1 for each population of adult minnows (*Phoxinus phoxinus*) at the onset of the experiment. Error bars represent ± 1 SE.

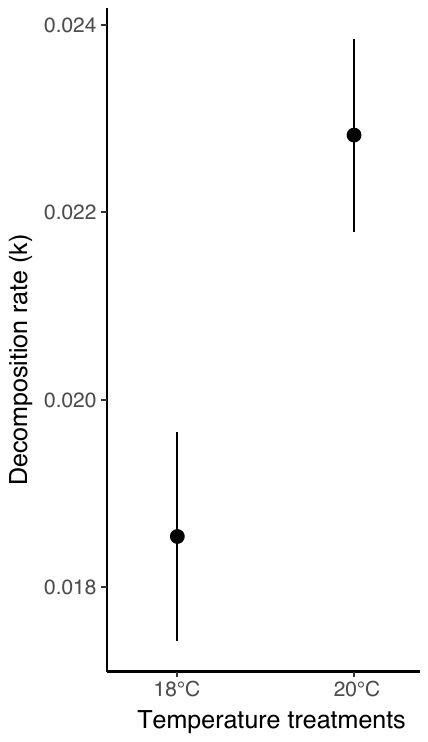

**Fig. S8.** Decomposition rate (k) at the end of the phase 1 for each temperature treatment. Error bars represent ± 1 SE.

**Table S1.** Results of the linear model analysis used to calculate effect size of intraspecific variation (among populations) and temperature on ecological and performance parameters.

|  |  | Temperature | | | Population | | | Interaction | | |
| --- | --- | --- | --- | --- | --- | --- | --- | --- | --- | --- |
|  |  | d.f. | F value | P-value | d.f. | F value | P-value | d.f. | F value | P-value |
| **Ecological parameters** | Cladocera abundance | 1 | 2.781 | 0.101 | 5 | 3.223 | 0.012 | - | - | - |
|  | Copepoda abundance | 1 | 0.005 | 0.942 | 5 | 0.703 | 0.623 | - | - | - |
|  | Size Copepoda | 1 | 2.298 | 0.135 | 5 | 2.053 | 0.084 | - | - | - |
|  | Size Cladocera | 1 | 1.261 | 0.266 | 5 | 2.360 | 0.051 | - | - | - |
|  | Bivalvia abundance | 1 | 14.437 | < 0.001 | 5 | 1.625 | 0.168 | - | - | - |
|  | Gastropoda abundance | 1 | 0.022 | 0.884 | 5 | 1.176 | 0.332 | - | - | - |
|  | Decomposition rate | 1 | 33.844 | < 0.001 | 5 | 1.534 | 0.194 | - | - | - |
|  | Pelagic algae stock | 1 | 0.117 | 0.734 | 5 | 0.867 | 0.509 | - | - | - |
|  | Benthic algae stock | 1 | 1.982 | 0.165 | 5 | 1.156 | 0.343 | 5 | 2.900 | 0.022 |
|  | Filamentous algae | 1 | 10.969 | 0.001 | 5 | 0.368 | 0.868 | - | - | - |
|  | Abiotic parameters | 1 | 0.002 | 0.968 | 5 | 0.852 | 0.519 | - | - | - |
| **Performance parameters** | Survival | 1 | 3.608 | 0.063 | 5 | 0.640 | 0.670 | - | - | - |
|  | Growth | 1 | 3.313 | 0.074 | 5 | 0.653 | 0.660 | - | - | - |
|  | Body condition | 1 | 0.089 | 0.766 | 5 | 0.537 | 0.747 | - | - | - |

**Table S2.** The causal pathways between variations in intraspecific, temperature and ecological parameters were obtained from path-analysis. Each response variable is directly or indirectly related to one or several independent variables. Independent variables that support an indirect effect are indicated by a subscript (“t+i” when the independent variable was directly related to both temperature and intraspecific variation, “i” when the independent variable was directly related only to intraspecific variation).

| Response variable | Independent variable | Path-coefficient | S.E. | *p*-value |
| --- | --- | --- | --- | --- |
| Abiotic parameters | Activity | -0.135 | 0.084 | 0.107 |
|  | Temperature | 0.219 | 0.089 | 0.014 |
|  | Filamentous algae^t+i^ | -0.757 | 0.101 | <0.001 |
|  | Pelagic algae stock^i^ | 0.303 | 0.079 | <0.001 |
| Bivalvia abundance | Activity | -0.172 | 0.105 | 0.102 |
|  | Temperature | 0.554 | 0.106 | <0.001 |
| Cladocera abundance | Body mass | 0.316 | 0.113 | 0.005 |
|  | Metabolic rate | 0.158 | 0.112 | 0.161 |
|  | Copepoda abundance^t+i^ | 0.333 | 0.102 | 0.001 |
| Size Cladocera | Bivalvia abundance^t+i^ | 0.367 | 0.123 | 0.003 |
| Copepoda abundance | Body mass | 0.615 | 0.162 | <0.001 |
|  | Growth rate | 0.533 | 0.176 | 0.003 |
|  | Temperature | -0.259 | 0.135 | 0.055 |
|  | Bivalvia abundance^t+i^ | 0.408 | 0.135 | 0.003 |
| Size Copepoda | Gut length | -0.28 | 0.126 | 0.027 |
| Decomposition rate | Aspect ratio | -0.337 | 0.115 | 0.004 |
|  | Body mass | 0.413 | 0.115 | <0.001 |
|  | Gut length | 0.171 | 0.105 | 0.103 |
|  | Metabolic rate | 0.234 | 0.124 | 0.059 |
|  | Temperature | 0.482 | 0.105 | <0.001 |
|  | Gastropoda abundance^i^ | 0.178 | 0.112 | 0.114 |
| Filamentous algae | Activity | 0.348 | 0.09 | <0.001 |
|  | Gut length | 0.164 | 0.09 | 0.07 |
|  | Metabolic rate | -0.25 | 0.09 | 0.005 |
|  | Temperature | 0.454 | 0.09 | <0.001 |
| Gastropoda abundance | Activity | -0.287 | 0.114 | 0.012 |
|  | Aspect ratio | 0.349 | 0.119 | 0.003 |
|  | Metabolic rate | -0.257 | 0.118 | 0.029 |
| Benthic algae stock | Body mass | -0.449 | 0.155 | 0.004 |
|  | Growth rate | -0.548 | 0.168 | 0.001 |
|  | Gut length | 0.191 | 0.108 | 0.076 |
|  | Temperature | 0.496 | 0.129 | <0.001 |
|  | Bivalvia abundance^t+i^ | -0.298 | 0.129 | 0.021 |
|  | Pelagic algae stock^i^ | 0.177 | 0.111 | 0.11 |
| Pelagic algae stock | Aspect ratio | -0.215 | 0.123 | 0.08 |
| Covariances | | | | |
| Body mass | Growth rate | -0.705 | 0.155 | <0.001 |
| Body mass | Metabolic rate | -0.458 | 0.142 | 0.001 |
| Growth rate | Metabolic rate | 0.344 | 0.127 | 0.007 |
| Metabolic rate | Aspect ratio | 0.241 | 0.12 | 0.045 |

**Table S3.** The causal pathways between variations in ecological parameters, temperature and performance parameters were obtained from path-analysis. Each independent variable that was related to temperature or intraspecific variation, and supports an indirect effect are indicated by a subscript (“t+i” when the independent variable was directly related to both temperature and intraspecific variation, “i” when the independent variable was directly related only to intraspecific variation, and “t” when the independent variable was directly related only to temperature variation).

| Response variable | Independent variable | Path-coefficient | S.E. | *p*-value |
| --- | --- | --- | --- | --- |
| Juvenile survival | Temperature | -0.245 | 0.128 | 0.055 |
|  | Benthic algae stock^t+i^ | 0.186 | 0.122 | 0.129 |
|  | Gastropoda abundance^i^ | -0.173 | 0.117 | 0.141 |
|  | Size Cladocera | -0.174 | 0.119 | 0.144 |
| Juvenile growth | Juvenile survival^t^ | -0.603 | 0.107 | <0.001 |
|  | Abiotic parameters^t+i^ | -0.239 | 0.107 | 0.026 |
|  | Gastropoda abundance^i^ | -0.201 | 0.104 | 0.054 |
| Juvenile body condition | Juvenile survival^t^ | 0.691 | 0.08 | <0.001 |
|  | Juvenile growth | 1.016 | 0.077 | <0.001 |
|  | Temperature | 0.282 | 0.086 | 0.001 |
|  | Abiotic parameters^t+i^ | 0.114 | 0.069 | 0.098 |
|  | Benthic algae stock^t+i^ | -0.115 | 0.069 | 0.098 |
|  | Bivalvia abundance^t+i^ | -0.311 | 0.082 | <0.001 |
|  | Copepoda abundance^t+i^ | 0.097 | 0.066 | 0.144 |
