## supplementary method for "Intraspecific variation alters ecosystem and next-generation performance as much as temperature"

*Community and ecosystem parameters measured at the end of the phase 1.*

(*i*) Pelagic algae stock was assessed as a proxy of pelagic primary productivity. Measurements were performed using a portable spectrometer (AlgaeTorch, bbe Moldaenke®) to assess the chlorophyll-a concentration (μg/L) in the water column. Two measurements were taken in each mesocosm and were averaged for the analyses.

(*ii*) Benthic algae stock was assessed as a proxy of the benthic primary productivity using a portable spectrometer (BenthoTorch, bbe Moldaenke®). The chlorophyll-a concentration (μg/cm^2^) was measured on two tiles (20 x 20 cm) placed in the mesocosms the day before the start of the experiment. These measurements were averaged for analyses.

(*iii*) The abundance of filamentous algae was quantified. Filamentous algae cover (%) was visually estimated by two operators, and values were averaged for analyses.

(*iv*) Zooplankton community was assessed by filtering 5 L of water through a 200 μm sieve. Samples were conserved in a 70% ethanol solution and subsequently identified to the order or family levels, including Copepoda (i.e., Cyclopoida and Calanoida) and Cladocera (i.e., Daphniidae, Chydoridae and Bosminidae). Zooplankton size was assessed by measuring 10 individuals of each order and family level from each mesocosm to the nearest 0.001 mm using ImageJ®.

(*v*) Decomposition rate was measured by quantifying the mass loss of black poplar (*Populus nigra*, a dominant riparian tree in southern France) abscised leaves(Alp et al. 2016). One day before the start of the experiment, 4 g of air-dried leaves were put in each mesocosm within a coarse mesh (1 x 1 cm) bag. At the end of the phase 1, the remaining leaf material was removed from the mesocosms, rinsed with tap water, oven dried at 60°C for three days and weighed to the nearest 0.001 g to assess the loss of biomass. The decomposition rate was calculated as $k=-\frac{ln\left( X \right)}{t}$ (Alp et al. 2016), where *X* is the proportion of litter remaining after phase 1 and *t* is the elapsed time in days.

(*vi*) Macroinvertebrates (> 1 mm, essentially molluscs) were collected from the mesh bags used to measure decomposition rates, conserved in a 70% ethanol solution, and identified as Bivalvia or Gastropoda.

(*vii*) Abiotic parameters of the water [pH, specific conductance (μS), oxygen concentration (mg.L^-1^) and turbidity (NTU)] were measured with a multiparameter probe (YSI Pro DSS Water Quality Meter®). We summarized these parameters using principal component axis (PCA) (package ade4 in R, Chessel et al. 2007)). We selected the first axis of the PCA as the synthetic variable. This axis explained 60% of the variance and was correlated to the oxygen concentration (loading component: -0.95), pH (-0.93), specific conductance (0.70) and, to a lesser extent, turbidity (0.25).

*Phenotypic measurement*

We measured five phenotypic traits on each adult fish in addition to the body mass to test for the contribution of each of these traits to ecosystem properties and transgenerational carry-over effects. We focused on traits that are likely to mechanistically affect ecosystem properties. Activity was measured throughout the first step of the experiment. Each morning the experimenters (AR and LDG) noted whether the fish were visible or not in the tank, and we considered a fish as active when it was visible. We then calculated the proportion of time each fish was visible to obtain an activity score. The growth rate (%.day^-1^) of the adults was calculated as the specific growth rate (SGR):$SGR=\frac{ln\left( Wf \right)-ln\left( Wi \right)}{T}*100$, where *Wf* and *Wi* are the final (after the experiment) and initial body masses, respectively, and *T* is time interval between two measurements (in days). To assess the metabolic rate we followed the procedure described in (Raffard et al. 2019). Briefly, each fish was individually placed in a metabolic chamber filled with 500 mL of dechlorinated tap water. Measurements of oxygen concentration were taken after 10 min, allowing individuals to acclimate, and continuously every five seconds for 50 min with oxygen probes. Chambers were set in a thermoregulated room at 17°C in the dark to lower the stress level. After one hour, fish were gently released in their home tank. Before to start the measurement, the individuals were starved for two days to ensure the same starvation level among individuals. Afterward fish were euthanized in a solution of benzocaine at 25 mg.L^-1^ to take four morpho-anatomical measurement. Body length (*Bl*), caudal fin depth (*Cfd*) and caudal fin surface (*Cfs*) were measured using picture analysis (ImageJ); gut length (*Gl*) was measured following dissection. We then calculated gut length as $\frac{Gl}{Bl}$ , and aspect ratio as $\frac{{CFd}^{2}}{CFs}.$

Alp, M., J. Cucherousset, M. Buoro, and A. Lecerf. 2016. Phenological response of a key ecosystem function to biological invasion. Ecology Letters 19:519–527.

Chessel, D., A. B. Dufour, and S. Dray. 2007. ade4: Analysis of ecological data: exploratory and euclidean methods in multivariate data analysis and graphical display. R package version:1–4.

Raffard, A., J. Cucherousset, J. G. Prunier, G. Loot, F. Santoul, and S. Blanchet. 2019. Variability of functional traits and their syndromes in a freshwater fish species ( *Phoxinus phoxinus* ): The role of adaptive and nonadaptive processes. Ecology and Evolution 9:2833–2846.
